# Enhancement of gut-microbiome-based classification of prodromal Parkinson’s Disease cases by hyperparameter tuning, integration of multimodal data, and meta-analysis

**DOI:** 10.64898/2026.10.18.759928

**Authors:** Shayamolima Gogoi, Pramod R Somvanshi, Vivek Thakur

**Affiliations:** Dept of Systems and Computational Biology, School of Life Sciences, University of Hyderabad, Gachibowli, Hyderabad, Telangana, India

**Keywords:** gut microbiome, Prodromal PD, RBD, Machine Learning, Binary Classifier, Random Forest Classifier

## Abstract

Parkinson’s Disease (PD) is often preceded by a prodromal phase marked by non-motor symptoms such as REM Sleep Behaviour Disorder (RBD), gastrointestinal dysfunction, and olfactory impairment. Mounting evidence implicates gut microbiota in the early pathophysiology of PD. In this study, we aimed to enhance the classification of prodromal PD using gut microbiota profiles, with a focus on model performance and biomarker identification.

Publicly available gut microbiome datasets of individuals with RBD and healthy controls were analyzed. Following batch correction and genus-level filtering, we identified six taxa significantly depleted in both RBD and PD groups— all belonging to short-chain fatty acid (SCFA)-producing genera.

Hyperparameter tuning on the testing set achieved an AUC of 0.889. When combining microbial and clinical features, the model achieved a peak mean AUC of 0.917. Feature selection identified 11 top-ranked features, yielding an AUC of 0.849. Using only the six differentially abundant SCFA-producing taxa, the model retained a strong predictive capacity with a mean AUC of 0.853.

These results demonstrate the utility of gut microbiota, particularly SCFA-producing taxa, in the early identification of prodromal PD. Integration with select clinical features further improves model performance, supporting their potential role in early diagnostic frameworks.

## Introduction

### Background and Rationale

Parkinson’s Disease (PD) is a progressive neurodegenerative disorder, first comprehensively described in 1817 by British physician James Parkinson in his seminal work “An Essay on the Shaking Palsy” (Parkinson, 1817). In this monograph, he documented the characteristic motor symptoms of PD, namely, tremor (shaking), rigidity (muscle stiffness), bradykinesia (slowness of movement), and postural instability (Parkinson, 1817; Lewis, 2012).

PD is one of the most common neurodegenerative diseases with over 8.5 million reported cases worldwide. The average age of PD onset is 60 years; however in recent times, there has been an increase in patients with disease incidences before the age of 50. PD incidence before the average age of incidence is called early-onset PD (Mehanna, 2022).

While the exact cause of PD is unknown, it’s believed to involve a combination of genetic and environmental factors, with the progressive loss of dopaminergic neurons in the brain, particularly in the substantia nigra, being the most prominent marker (Steece-Collier et al., 2002). The pathological hallmark of PD is the presence of Lewy bodies - aggregated clumps of α-synuclein in the brain (Spillantini, M., 1997).

The phase that precedes the irreversible motor symptoms of PD is known as the prodromal phase. A wide range of non-motor symptoms are observed in this phase, such as constipation, hyposmia, olfactory and cognitive impairment, and REM-sleep behavior disorder (RBD) (Berg et. al., 2015). The Movement Disorder Society (MDS) Research Criteria for Prodromal PD provide an evidence-based methodological framework for statistically estimating the likelihood of prodromal PD in an individual, and among multiple factors, RBD has the highest likelihood (Heinzel et al., 2019). Rapid Eye Movement (REM) sleep behavior disorder is a parasomnia characterized by abnormal movements during REM sleep. One of the most common symptoms of RBD is the enactment of dreams while in REM sleep. RBD is diagnosed by polysomnography according to the International Classification of Sleep Disorders (ICSD). Three types of RBD are known: idiopathic (iRBD), drug-induced, and secondary RBD due to medical conditions (Högl et. al., 2018) (American Academy of Sleep Medicine, 2023). iRBD is most suggestive in neurodegenerative diseases like dementia with Lewy Bodies (DLB) and Parkinson’s disease (Postuma et. al., 2019). In fact, it is estimated that 50-70% of individuals diagnosed with RBD go on to develop PD (Pham, Sankari, & Slowik, 2024).

### Current Status and gaps

To date, only five studies have examined gut dysbiosis in RBD, a prodromal stage of PD, summarized in Table 1. Among them, only Huang et al. (2023) attempted to harness this information for predictive modeling. Using a Random Forest Classifier based on 12 genera, the study achieved moderate performance (mean Area Under Curve, AUC = 0.75) on an internal test set (Figure S1).

**Table 1:** Overview of the studies reporting gut dysbiosis in RBD.

| Study | Country | Sample*<br>Distribution | Status of microbes<br>(RBD with respect to HC*) |  |
| --- | --- | --- | --- | --- |
|  |  |  | Enriched | Depleted |
| **PRJEB52086**<br>(Huang, et al., 2023) | China<br>(Hong Kong) | HC = 108<br>FDR = 127<br>RBD = 170<br>PD = 36 | *Akkermansia*,<br>*Desulfovibrio* | *Roseburia*,<br>*Lachnospiraceae*<br>*ND3007_group*,<br>*Faecalibacterium*,<br>*Butyricicoccus* |
| **PRJDB9049**<br>(Nishiwaki, et al., 2020) | Japan | HC = 137<br>RBD = 26<br>PD = 223 | *Rikenellaceae*<br>*Akkermansiaceae* | *Ruminococcus*,<br>*Alistipes*, *Akkermansia*,<br>[*Eubacterium*]<br>*coprostanoligenes group*,<br>*Family XIII AD3011 group* |
| **PRJNA822715**<br>(Zhang, et al., 2024) | China | HC = 60<br>*i*RBD = 35<br>PD = 94 | *Aerococcus*,<br>*Eubacterium*,<br>*Gordonibacter*,<br>*Stenotrophomonas* | *Butyricicoccus*,<br>*Faecalibacterium*,<br>*Haemophilus* |
| **PRJNA381395**<br>(Heintz-Buschar, et al., 2017) | Germany | HC = 78<br>*i*RBD = 21<br>PD= 76 | *Akkermansia* | *Prevotellaceae* |
| **(Du, et al., 2024)** | China | HC = 35<br>*i*RBD = 29<br>MSA = 42<br>PD = 40 | NA | *Lachnospira*, *Veillonella*,<br>*Bacteroides*,<br>*Faecalibacterium* |

\*HC – Healthy Control; RBD – REM Sleep Behavior Disorder; *i*RBD – Idiopathic RBD; MSA – Multiple System Atrophy

In terms of data integration and scalability, no existing study has attempted to combine clinical and microbial features into a unified predictive model for RBD. The TREND study (Tübingen Evaluation of Risk Factors for Early Detection of Neurodegeneration) aimed to identify correlations among established risk and prodromal markers of PD and gut microbial composition in an elderly population enriched for PD risk (Heinzel et al., 2021). This study found that markers such as physical inactivity, occupational solvent exposure, and constipation were significantly associated with α-diversity, whereas physical inactivity, biological sex, constipation, possible RBD, and smoking were significantly associated with β-diversity (Heinzel et al., 2021). This shows that integrating multimodal data could enhance both predictive performance and biological interpretability, which remain clear gaps in the field.

Attempts at meta-analysis are also sparse. While Nishiwaki et al. (2019) combined their Japanese cohort with a German dataset (Heintz-Buschart et al., 2017), the limited sample size made their findings inconclusive. Notably, their analysis did not observe a significant reduction in SCFA-producing bacteria in RBD, a hallmark of PD-related dysbiosis, further complicating the gut-brain axis narrative in prodromal stages. A more recent preprint (Arıkan, 2025) expanded the scope by pooling four datasets (n=973; 343 HC, 251 RBD, 379 PD) and applying a LASSO classifier. However, their mean 10-fold cross-validation (CV) AUC was modest (0.61 ± 0.10), reflecting both cross-cohort variability and incomplete feature utilization.

### Hypothesis and overview of the work

Given the gaps in the performance of the classification studies mentioned above, we explored options to enhance model performance using the existing data. Among the strategies explored are hyperparameter tuning, integration of multimodal data, and increasing the sample size through meta-analysis. The hypotheses underlying them are explained below:

#### Hyperparameter tuning to improve baseline model performance

The performance of existing models (binary random forest classifier) can be improved further by systematically tuning hyperparameters rather than relying on the default or tested settings. We revisited the existing Random Forest-based classifier in Huang et. al. (2023) and implemented grid hyperparameter tuning, optimizing parameters such as the number of estimators, tree depth, and feature sampling strategy using bootstrapping and ROC-AUC scoring. This step served as a foundation for model refinement.

#### Integration of available clinical features with the taxonomic features

Incorporating easily available clinical metadata alongside taxonomic features may improve classification performance, reflecting the multifactorial nature of prodromal PD. We constructed combined feature matrices that included both Centred Log Ratio (CLR)-transformed taxonomic abundances and normalized clinical variables, and trained the model on this integrated dataset. Comparative evaluation was done against microbiome-only models using metrics such as ROC-AUC, balanced accuracy, and precision-recall curves.

#### Meta-Analysis using an expanded dataset across cohorts

Given a small effect size of taxonomic and/or basic clinical features, increasing sample size and diversity by including independent datasets may enhance model generalizability and allow for the detection of robust, cohort-invariant microbial signatures. We curated and harmonized an additional dataset with a similar subject recruitment strategy. After appropriate batch effect correction, the datasets were merged to build a larger, meta-analyzed training set. We then reassessed model performance and feature importance to identify consistently predictive taxa.

## Materials and methods

An overview of the steps and datasets involved in the study is shown as a flowchart in Figure S2.

### Selection of Datasets

Relevant publications were identified in PubMed using the search terms: (“Microbe” OR “Microbiota” OR “Microbiome”) AND (“RBD” OR “REM”) AND (“Parkinson” OR “PD”).

The selection criteria for inclusion in the study were: (i) No reviews or systematic reviews, (ii) Extraction of DNA from human stool samples, (iii) 16S rRNA amplicon sequencing. The datasets were downloaded from NCBI.

### Hyperparameter Tuning

The dataset from Huang et al. (2023) (NCBI Bioproject ID: PRJEB52086; see Table 1 for details about the groups), which included Centred Log Ratio (CLR)-transformed abundances of selected microbial genera along with clinical metadata, was used for hyperparameter optimization. An 80:20 split was applied to create training and test sets. GridSearchCV from scikit-learn was employed on the training data to tune hyperparameters of the Random Forest Classifier (Pedregosa et al., 2011; Gilbert et al., 2023). The optimized hyperparameters are explained below:

a. *bootstrap = True*: This means each decision tree is trained on a random sample (with replacement) from the training data. It helps introduce variability and prevents overfitting by ensuring each tree sees a slightly different version of the data.
b. *max_depth = 5*: Limits the depth of each tree to level 5. Shallow trees reduce the risk of overfitting and force the model to make more general decisions rather than memorizing the data.
c. *max_features = ‘sqrt’*: At each split, the algorithm considers only the square root of the total number of features. This randomness increases model diversity.
d. *min_samples_leaf = 1*: A leaf node must have at least one sample. Lower values allow the model to fit fine details.
e. *min_samples_split = 5*: A node must have at least 5 samples to be considered for a split. This prevents overly small splits that could lead to noise fitting.
f. *n_estimators = 200*: The model builds 200 trees and combines their outputs, thereby providing better performance and robustness.

### Integration of clinical and taxonomic features

Metadata corresponding to the above study (PRJEB52086, Huang et al., 2023) was downloaded and curated. Among the available clinical features, their selection was based on their likelihood of prodromal PD and clinical accessibility. Only samples that contained complete data for all of these selected features were retained for further analysis. For classification, three feature sets were evaluated:

a. **Microbial-only**: Centred Log Ratio-transformed (CLR-transformed) abundances of twelve selected genera.
b. **Clinical-only**: The six selected clinical features.
c. **Integrated**: A combined set including both the twelve taxa and six clinical features.

For each feature set, model hyperparameters were optimized using GridSearchCV with 5-fold cross-validation.

### Meta-Analysis

#### a. Preprocessing the datasets

Out of four publicly available gut microbiome datasets (Table 1), only two were selected for analysis based on sequencing strategies and subject recruitment criteria (NCBI BioProject IDs: PRJEB52086 and PRJDB9049).

Each dataset was pre-processed as follows: first, the raw sequencing reads were downloaded from NCBI SRA. The quality assessment was performed using FastQC (Andrews, S., 2010), with aggregate reporting via MultiQC (Ewels, Magnusson, Lundin, & Käller, 2016). The quality filtering and adapter trimming were carried out using *fastp* (Chen, 2023). The filtered reads were processed in QIIME2-amplicon-2024.10 (Boleyn, Rideout, Dillon, Bokulich, & Caporaso, 2019), including demultiplexing, denoising, and feature table construction. Taxonomic classification was performed using the SILVAv138.2 database (Quast, 2013; Yilmaz, 2014). Amplicon sequence variants (ASVs) associated with mitochondria and chloroplasts were removed to exclude non-bacterial sequences. The ASV count table was collapsed to the genus level. Prevalence and abundance filtering was applied - only genera present in at least 10% of samples and with a minimum relative abundance of 0.05% were retained. These thresholds were selected as pragmatic preprocessing criteria to retain genera with sufficient prevalence and abundance for downstream analyses (Nearing et. al., 2022). The resulting genus-level feature table was exported using the *biomformat* (Version 1.34.0) package in R.

#### b. Batch Correction

Following feature filtering, batch correction was performed using the Microbiome Batch Effects Correction Suite (MBECS) (Olbrich, Künstner, & Busch, 2023) package in R to minimize dataset-specific variability and enhance comparability between the two selected studies.

Four batch correction algorithms were tested:

a. ComBat (bat): A popular empirical Bayes method that adjusts for batch effects by modeling both biological and technical variation. It assumes data are normally distributed and works well for large sample sizes.
b. Remove Batch Effect (rbe): A linear modeling approach implemented in the *limma* package. It directly removes the batch-associated variance from the data, preserving biological signal while eliminating technical noise.
c. Support Vector Decomposition (svd): A matrix decomposition technique that identifies and removes variation associated with batch labels. It is useful when batch effects are captured in the dominant components of variation.
d. Batch Mean Centering (bmc): A simpler method that adjusts each batch to have the same mean across features.

### Differential Abundance Analysis

Raw genus-level count tables were used to identify taxa that were differentially abundant between the study groups. Analysis was conducted using the ANCOM-BC package in R, which accounts for compositional and sampling biases (Lin, 2020). The Benjamini-Hochberg (BH) method was employed to correct for multiple testing and control the false discovery rate.

### Feature Selection

The combined dataset was split 80-20 into training and testing sets, respectively. A feature selection pipeline was run on the training set, incorporating four methods, namely Recursive Feature Elimination (RFE), Sequential Feature Selection Forward (SFS-F), Sequential Feature Selection Backward (SFS-B), and SHapley Additive exPlanations (SHAP) importance score (Lundberg & Lee, 2017). The ranks of the taxa were averaged across all four algorithms, and the minimum set of features that collectively achieved a significantly high AUC score on the training set was used to build a machine learning model.

### Model Building and Assessment

Random Forest Classifier was used to construct the model. Bootstrapping with 1000 iterations was performed, and the mean evaluation metrics were obtained and plotted using scikit-learn and matplotlib libraries in Python.

## Results

### Hyperparameter tuning

Using the same taxa-based features as reported in the earlier study involving Random Forest-based classification (Huang et al., 2023), a grid search was run to obtain the hyperparameters with the highest performance. The optimized hyperparameters were: ‘bootstrap’: TRUE, ‘max_depth’: 5, ‘max_features’: ‘sqrt’, ‘min_samples_leaf’: 1, ‘min_samples_split’: 5, ‘n_estimators’: 200. The newly trained model outperformed the model reported in Huang et. al. (2023) despite using the same features and dataset. A mean ROC-AUC of 0.889 was reported (Figure 1), as opposed to 0.75 in the previous study (Figure S1). The Receiver Operating Characteristic Curve – Area Under Curve (ROC-AUC), a metric that summarizes a model’s ability to discriminate between classes, ranges from 0.5 (random guessing) to 1 (perfect classification). An AUC of 0.889 indicates that the model can correctly distinguish between RBD and Control samples nearly 89% of the time.

**Figure 1:**
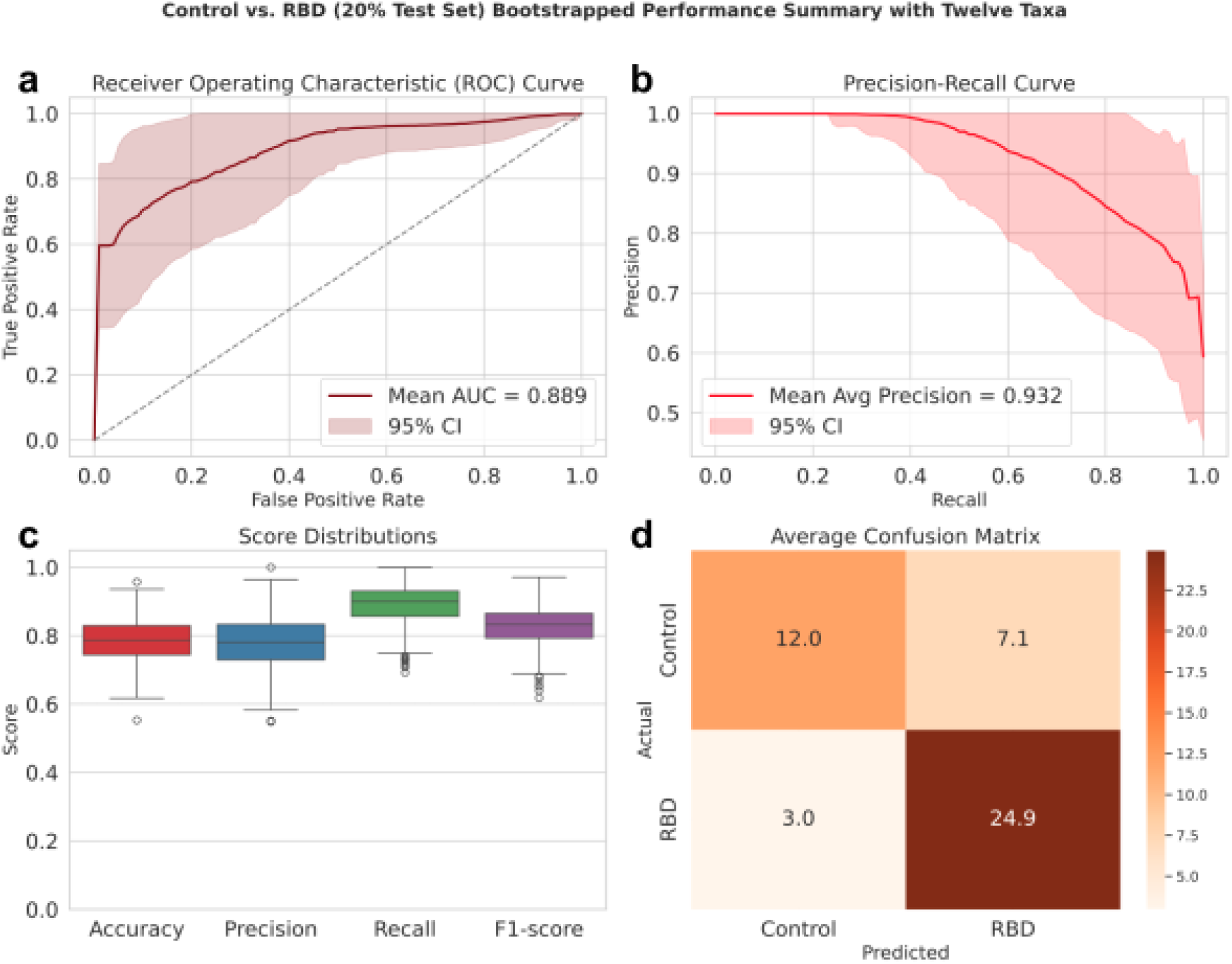
Random Forest Model for RBD with hyperparameter tuning. Performance of the tuned binary classifier on the 20% test set using the twelve original taxonomic features reported in the earlier study (Huang et al., 2023). a) The ROC curve showed a mean AUC of 0.889 on the test set across 1000 bootstrapping iterations. b) The Precision-Recall curve showed Mean Average Precision to be 0.93. c) The performance scores showed that the mean accuracy was 0.79, the mean precision was 0.78, the mean recall was 0.89, and the mean F1-score was 0.83. d) The confusion matrix.

### Integration of clinical and taxonomic features

In addition to hyperparameter tuning, it was examined whether including clinical features in the existing binary classification model, which had so far used only microbial features, would further improve the model (PRJEB52086, Huang et al., 2023). From the clinical features available in the metadata, six were selected: olfactory impairment, age, Bowel Movement Frequency Score (BMF score), Rome Incomplete Evacuation, Rome Straining, and BMI (see methods). Even using just the six clinical features, a mean ROC-AUC of 0.894 was reported on the testing set (Figure S3). On the combined set of clinical and taxonomic features, model performance was slightly improved, with a mean AUC of 0.914 (Figure 2); however, other model evaluation metrics did not show significant improvement (Figure S4). In order to know the relative contribution of the clinical and microbial features, they were ordered based on their correlation with the group (RBD), using the SHAP values, and the clinical features were often ranked ahead of the microbial ones (Figure 3), indicating the importance of using diverse feature categories for classification.

**Figure 2:**
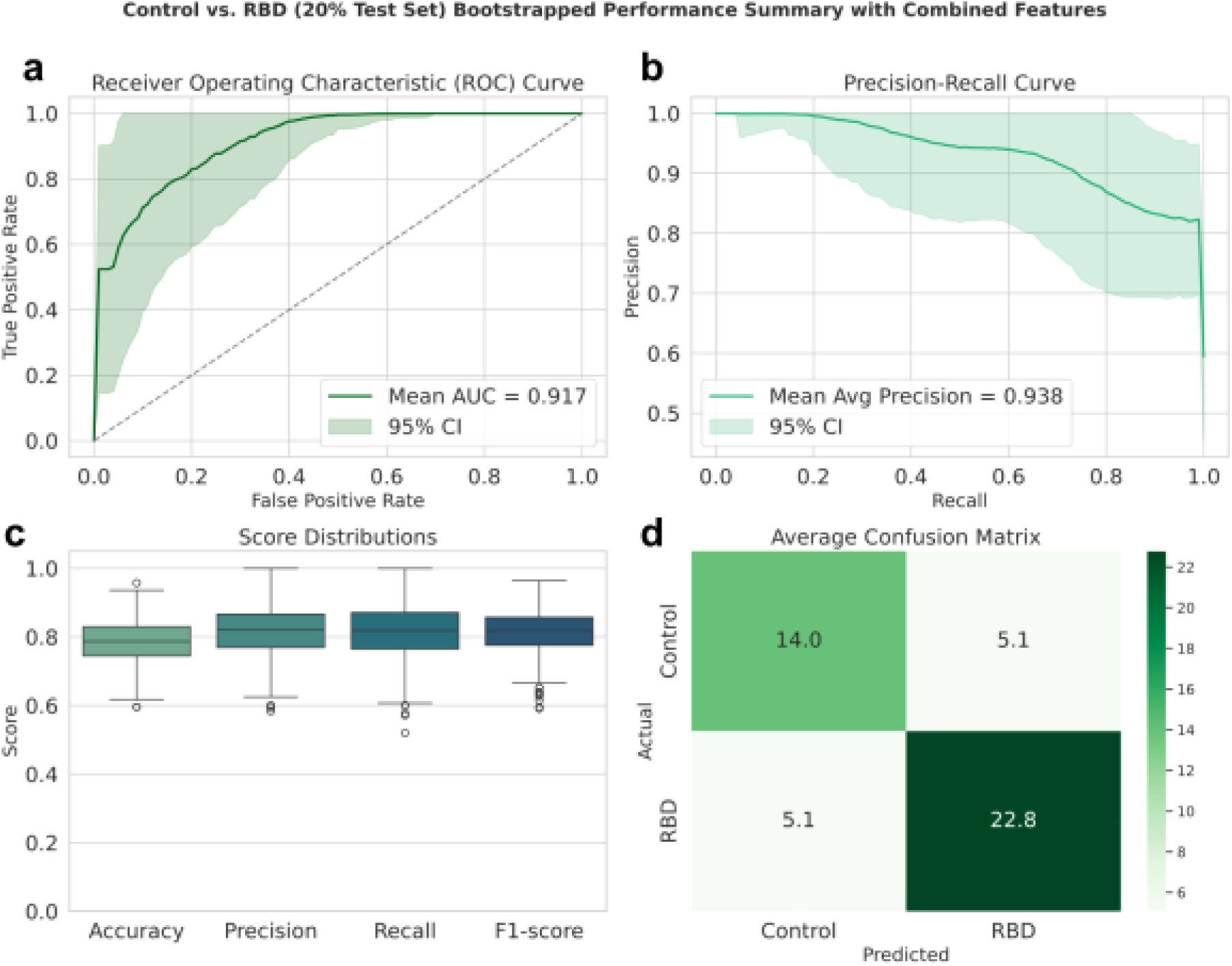
Binary classifier model with combined features. Performance of the tuned binary classifier on the 20% test set using the original taxonomic features and six clinical features. a) The ROC curve showed a mean AUC of 0.917 on the test set across 1000 bootstrapping iterations. b) The Precision-Recall curve showed Mean Average Precision to be 0.938. c) The performance scores showed that the mean accuracy was 0.79, the mean precision was 0.84, the mean recall was 0.83, and the mean F1-score was 0.83. d) The confusion matrix.

**Figure 3:**
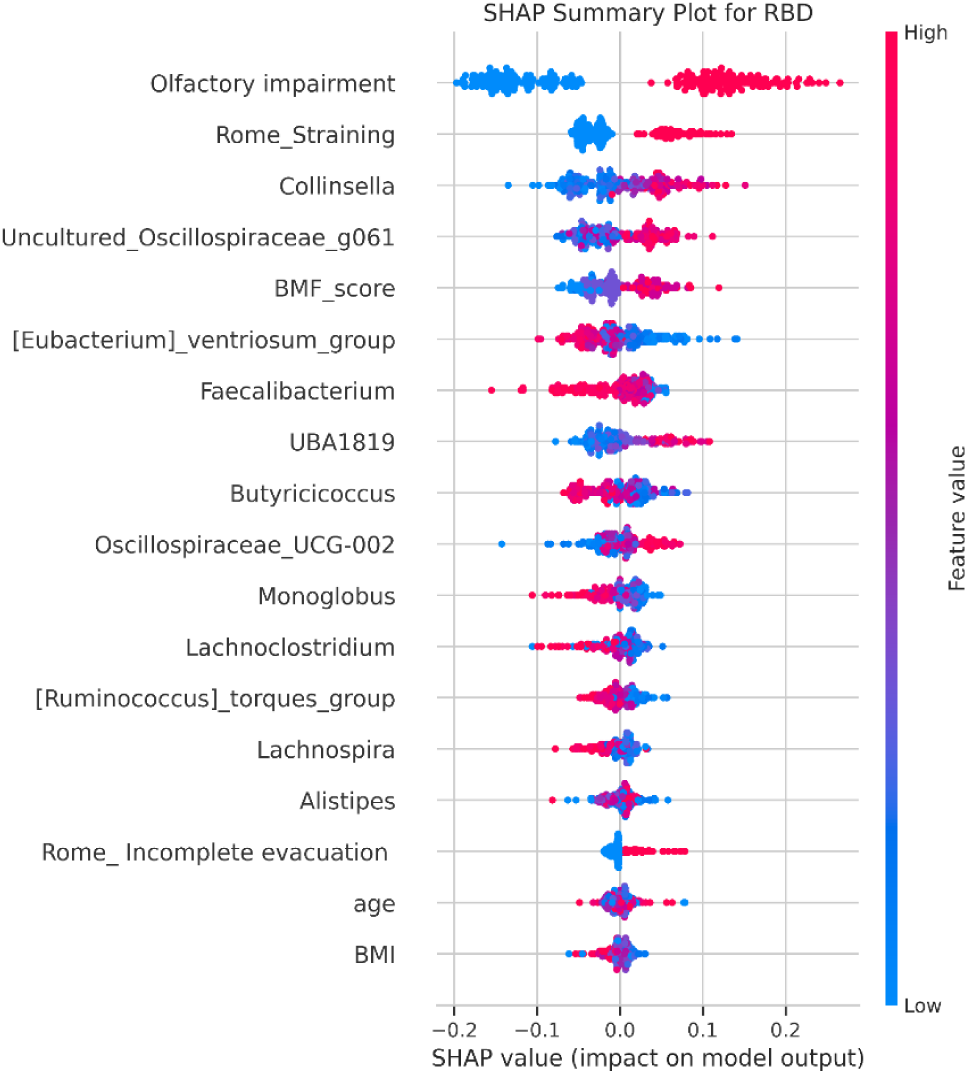
SHAP summary plot for RBD with combined features. The distribution of the integrated features across Control and RBD. SHAP value greater than 0 implies positive correlation with RBD.

### Meta Analysis: Sample selection & Batch Correction

Out of four datasets, two were found suitable for meta-analysis: PRJEB52086 (Huang et al., 2023) and PRJDB9049 (Nishiwaki et al., 2020) (see methods). Before merging the datasets for model training, it was important to assess and correct for batch effects, which can arise due to differences in sample processing or sequencing runs across datasets.

Out of four batch correction algorithms tested, ‘Remove Batch Effect’ (rbe) was found to be the most effective (Figure S5a,b), reducing the variance attributable to batch effects from 26.36% to 0%. This was visually confirmed by PCoA plots (Figure 4a,b), where batch-driven clustering observed before correction was no longer present post-correction, indicating successful harmonization of the datasets. Total number of samples after batch correction grew to: Control = 245, RBD = 196, PD = 259.

**Figure 4:**
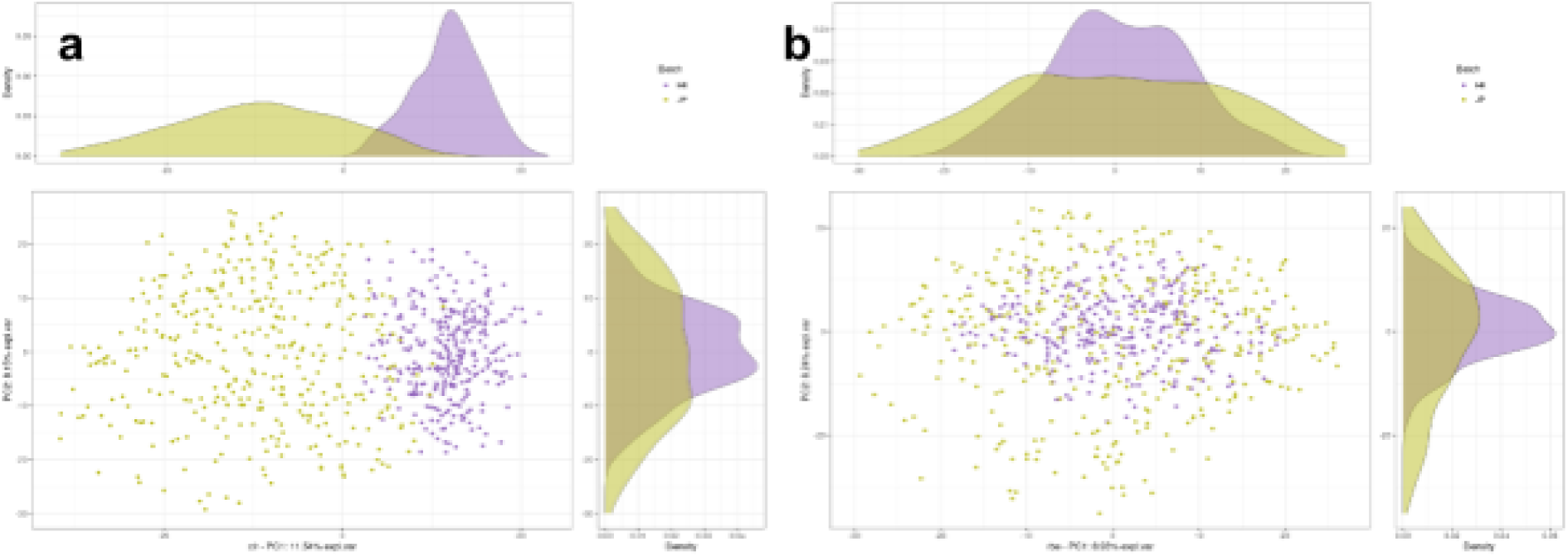
Principal Coordinate Axes (PCoA) plots depicting the effect of batch correction. a) Before batch effect correction, samples cluster distinctly by batch, indicating substantial technical variability across datasets. The first principal coordinate (PC1) accounts for 11.54% of the total variance, while the second (PC2) explains 8.15%. b) After applying the Remove Batch Effect (RBE) method, batch-associated clustering is no longer evident. The explained variance of PC1 reduces to 8.93%, and PC2 slightly increases to 8.24%, reflecting successful removal of batch-related variation while retaining biological signal integrity. HK - Hong Kong (Huang et al., 2023), JP - Japan (Nishiwaki et al., 2020).

### Meta-Analysis: Feature Selection

To identify the optimal number of features for model construction, the trade-off between the number of features and ROC-AUC was plotted. Although 28 features yielded the highest AUC score, even with 11 features, the score wasn’t very different (Figure 5a). Thus, the top eleven taxa, whose SHAP values are shown in Figure S6, were chosen for further downstream analyses and ML-model training.

**Figure 5:**
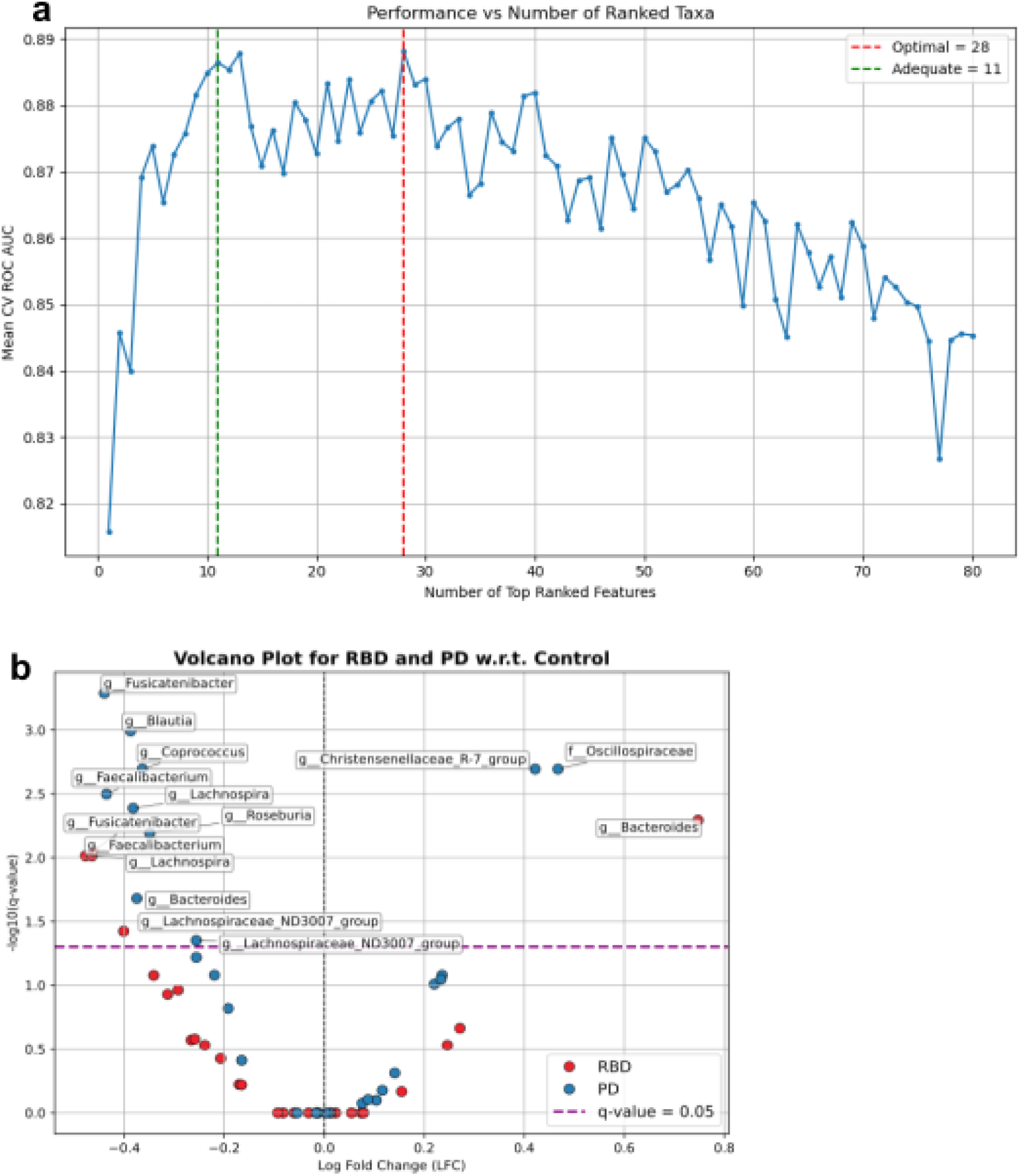
Feature selection using different strategies. **a)** Trade-off between performance and features. The X-axis plots the number of features in ascending order of their average rank. The Y-axis shows the mean 5-fold cross-validation (CV) ROC AUC score. **b)** Volcano plot of differentially abundant taxa in RBD and PD. Only six common taxa fall below the FDR threshold of 0.05.

As an alternative to feature selection, Differential abundance analysis was performed to identify microbial taxa that consistently distinguish RBD and PD subjects from healthy controls. A total of 28 taxa were found to be differentially abundant in both disease conditions across both cohorts, suggesting potential shared microbial signatures associated with neurodegenerative progression (Figure S7). On correcting for multiple comparisons using the false discovery rate (FDR), only 6 taxa remained statistically significant (*q* < 0.05), indicating high confidence in their association with disease states (Figure 5b). Notably, all six of these taxa are either known or predicted to be involved in short-chain fatty acid (SCFA) production (Table 2).

**Table 2:** Significantly differentially abundant taxa in RBD and PD (compared to Healthy Control subjects)

| Genus | Status in RBD | Status in PD | SCFA produced |
| --- | --- | --- | --- |
| *Fusicatenibacter* | Depleted | Depleted | Butyrate |
| *Blautia* | Depleted | Depleted | Acetate and butyrate |
| *Faecalibacterium* | Depleted | Depleted | Butyrate |
| *Lachnospira* | Depleted | Depleted | Butyrate |
| *Lachnospiraceae ND3007 group* | Depleted | Depleted | Likely butyrate |
| *Bacteroides* | Enriched | Depleted | Acetate/propionate |

The consistent depletion or enrichment of these SCFA-producing microbes in both RBD and PD reinforces the hypothesis of a gut microbial dysbiosis contributing to prodromal and overt Parkinson’s disease pathology. These taxa may serve as robust biomarkers and potential targets for therapeutic interventions aimed at restoring SCFA balance.

### Meta-Analysis: Model Training and Evaluation

To evaluate the predictive value of taxa identified through different analytical strategies, a Random Forest binary classifier was trained under two distinct conditions, using optimized hyperparameters obtained through grid search in each case:

i. Using the six differentially abundant (DA) taxa identified consistently in both RBD and PD groups.
ii. Using eleven taxa selected via a model-based feature selection pipeline, based on importance rankings.

In the DA-based model, the classifier achieved a mean ROC-AUC of 0.853, while the feature-selected model yielded almost a similar mean ROC-AUC of 0.849 (Figure 6a,b). Although the performance difference was modest, the DA taxa-led model showed superior results, suggesting that biologically meaningful features—particularly those consistently associated with disease states—offer a robust signal for classification (Figure 6c). This reinforces the importance of combining statistical significance with domain relevance in predictive model design.

**Figure 6:**
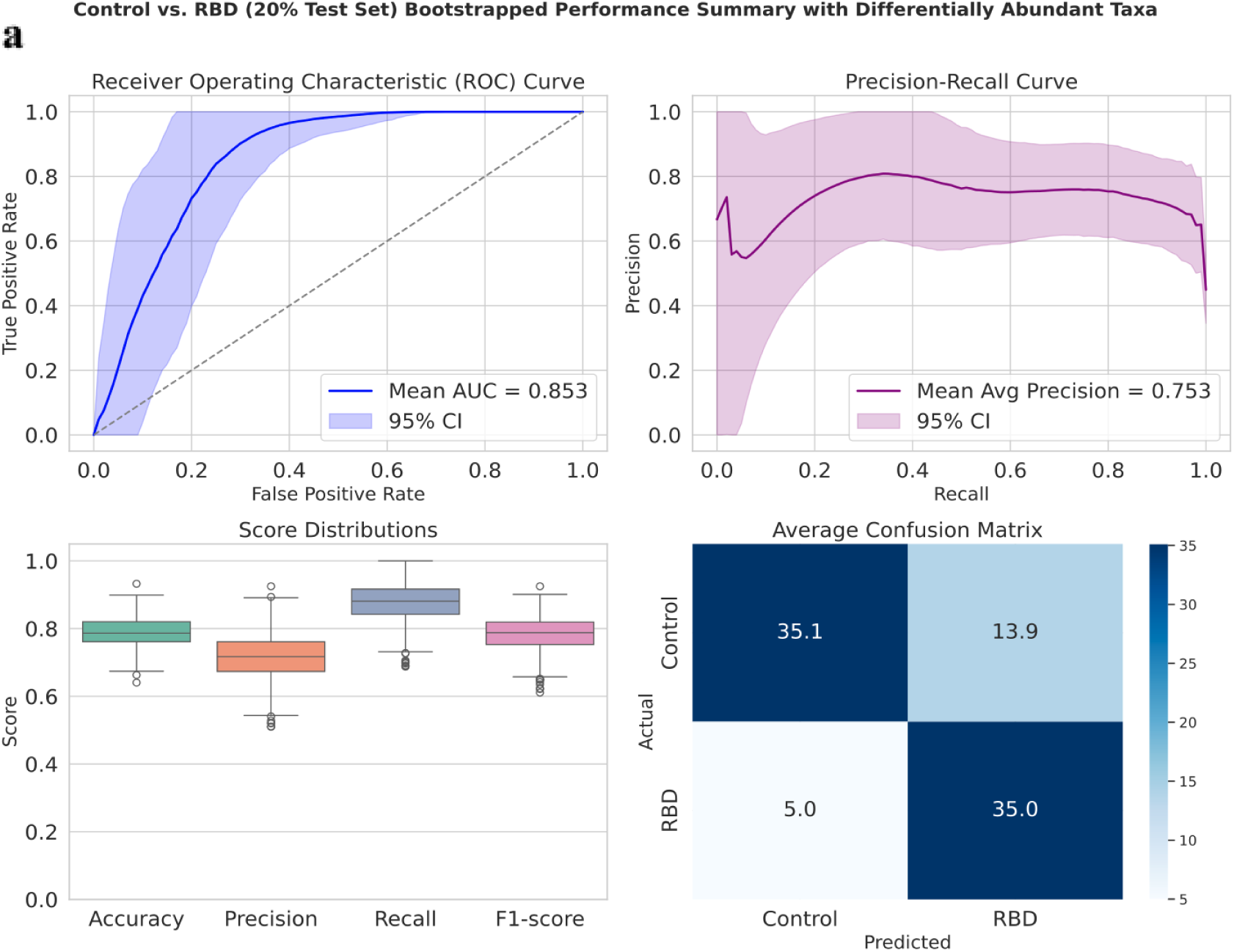
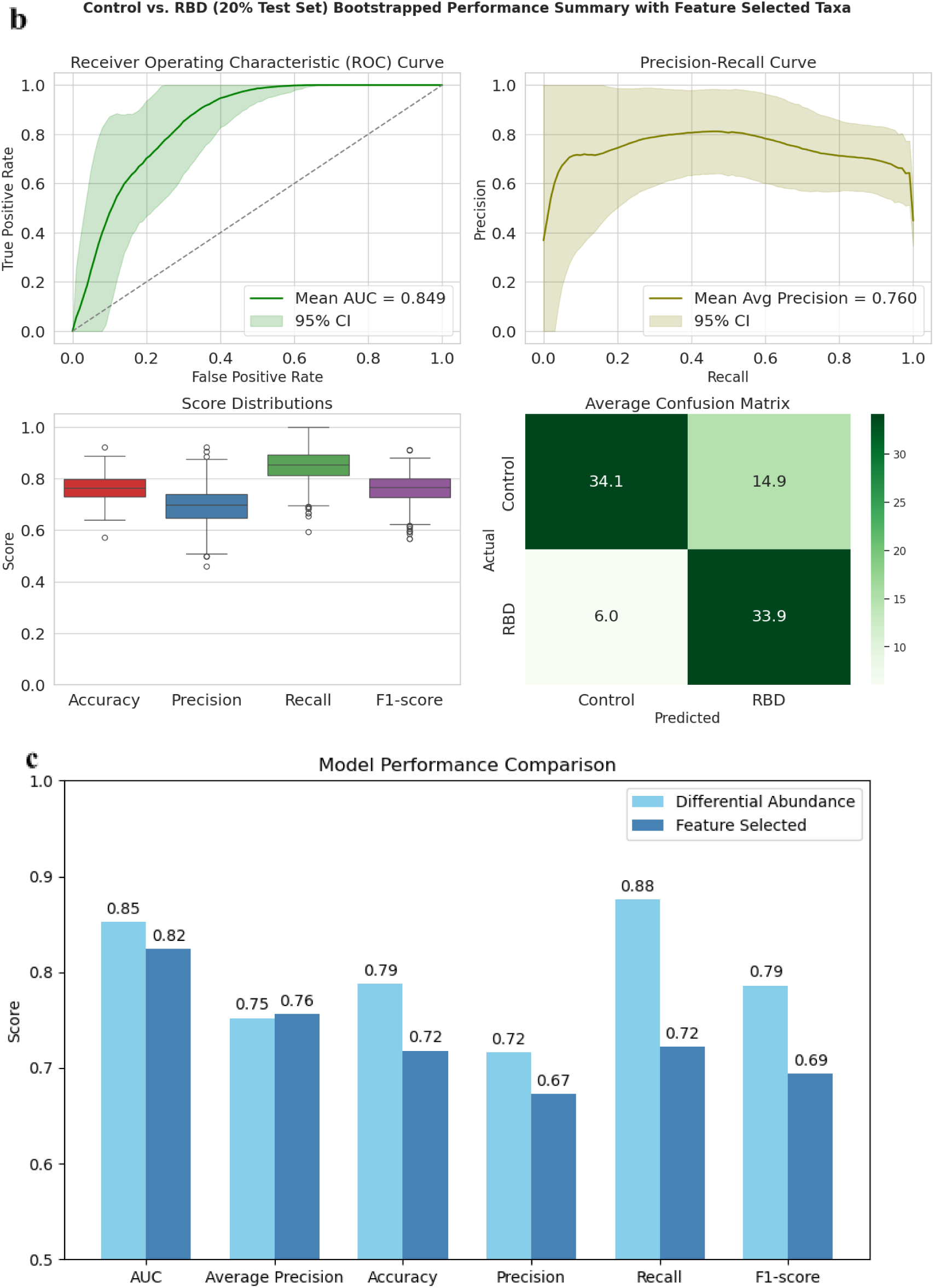
Meta-analysis: Random Forest Classifier Performance. a) Using six DA taxa. b) Using eleven feature-selected taxa. c) Model Performance Comparison (DA and Feature-Selected). Evaluation metrics of the model trained with differentially abundant taxa and the model trained with feature-selected taxa are plotted adjacent to each other. The dataset with differentially abundant taxa has consistently higher metric scores than feature-selected taxa.

Furthermore, when the six differentially abundant (DA) taxa were used to classify RBD versus PD, the model achieved a remarkably high mean ROC-AUC of 0.963, indicating excellent discriminative performance (Figure S8). This strong separation between the two groups suggests that these taxa capture key microbiome alterations associated with the transition from the prodromal to the clinically manifest stage of Parkinson’s disease.

Interestingly, the ability of a small set of microbial features to distinguish RBD from PD supports the hypothesis that gut dysbiosis may not be static, but instead follows a progressive trajectory as the disease advances. In this context, RBD — widely recognized as a prodromal phase of PD — may represent an intermediate state in a continuum of gut microbial shifts. The observed differences in microbial composition between RBD and PD imply that certain taxa either emerge, decline, or undergo compositional changes as neurodegeneration progresses.

## Discussion

The findings of this study underscore the importance of hyperparameter tuning in optimizing machine learning model performance, as observed in the case of the existing binary classifier for RBD and control cases, based on gut microbial features. Notably, even when using the same features and dataset, we observed significant improvements in classification accuracy following systematic optimization of model parameters. This highlights the importance of refining model architecture in addition to selecting informative features.

Integrating readily available clinical features with gut microbiome-derived taxonomic signatures demonstrated additional value in enhancing predictive performance. Although the observed gain was modest, the consistent trend suggests that a combined feature set provides a more comprehensive and biologically informed view of disease status. This supports the idea that leveraging multi-modal data could offer practical benefits for early detection and risk stratification in neurodegenerative disorders.

Intriguingly, our meta-analysis revealed a consistent depletion of short-chain fatty acid (SCFA)-producing microbes in both the RBD and PD groups, which contrasts with the findings of Nishiwaki et al. (2019), who reported a depletion of SCFA-producers only in PD. Such discrepancies may arise from differences in bioinformatics pipelines, taxonomic resolution, or statistical testing methods. While the depletion of SCFA-producing taxa in PD is well-established across multiple studies, their reduced abundance in RBD observed here supports the hypothesis of a progressive pattern of gut dysbiosis from the prodromal to clinical stages of Parkinson’s disease. This not only reinforces the concept of microbiome involvement in disease pathogenesis but also lays the groundwork for experimental studies evaluating the diagnostic and therapeutic potential of SCFA-producers in early-stage PD.

It is important to acknowledge the limitation that only two of the four publicly available RBD gut microbiome datasets were included in our meta-analysis. The remaining two were excluded due to limitations in their QIIME2-processed feature tables, which lacked sufficient genus-level taxonomic resolution. These issues may stem from low sequencing depth, poor-quality reads, or suboptimal data processing workflows. We also recognize the possibility of technical oversights on our part during preprocessing, and we report this limitation transparently.

Another key limitation lies in the geographic and ethnic homogeneity of the included cohorts, both of which originate from East Asian populations (China and Japan). Given the influence of diet, lifestyle, and genetics on gut microbiota composition, this homogeneity limits the generalizability of our findings to more diverse global populations. Furthermore, the lack of detailed metadata for the excluded datasets restricted our ability to assess the integrative predictive value of combining clinical and microbiome features on a broader scale.

In summary, this study highlights the potential of microbial and clinical markers in classifying prodromal and clinical stages of PD, supports a progressive model of gut dysbiosis, and calls for more standardized, high-quality, and globally representative datasets to advance microbiome-based diagnostics in neurodegeneration.

### Future Perspectives

Future work should focus on obtaining and analyzing high-quality and well-annotated RBD gut microbiome datasets. The inclusion of metadata – including clinical features, dietary habits, medication, and disease history-will allow deeper insights into consistent microbial signatures associated with prodromal PD. Secondly, investigating the gut microbiome of prodromal individuals in Indian cohorts will help uncover region-specific microbial patterns and signatures, which could be influenced by genetic, environmental, and dietary habits unique to the Indian subcontinent. Additionally, an important future direction is the design of a robust multiclass classification model capable of distinguishing between different stages of Parkinson’s Disease from the healthy:prodromal, early-stage, and late-stage PD. Such models could improve clinical decision-making and patient stratification for early interventions. Lastly, early-onset PD may follow a different pathophysiological trajectory compared to typical late-onset PD. Investigating the gut microbiome specific to EOPD cases could reveal unique biomarkers and provide insights into the clinical etiology and progression of this disease subgroup.

## Author contributions

Conceptualization: VT; Methodology: SG; Investigation: SG; Software and Scripts: SG; Formal Analysis: SG, VT, and PRS; Writing – Original Draft: SG; Writing – Review & Editing: VT; Supervision: VT; Funding Acquisition: VT. All authors reviewed and approved the final version.

## Acknowledgements

The authors would like to thank Dr. Anubha Dey for her input on feature selection, Mr. Valluru Devesh for generating preliminary data on this topic, and members of the VT lab (Nisha, Angeo, Sachidanand, Manas, and Suma) for their critical inputs on the work.

## Data availability

The publicly available gut microbiome sequencing datasets analyzed in this study were obtained from the NCBI Sequence Read Archive (SRA) under BioProject accession numbers PRJEB52086 and PRJDB9049. The codes used for data processing and analysis are available in the GitHub repository at: https://github.com/shaya03/Gut-microbiome-based-classification-of-prodromal-Parkinson-s-Disease

