## Supplementary Data for "Enhancement of gut-microbiome-based classification of prodromal Parkinson’s Disease cases by hyperparameter tuning, integration of multimodal data, and meta-analysis"

### Supplementary figures and tables

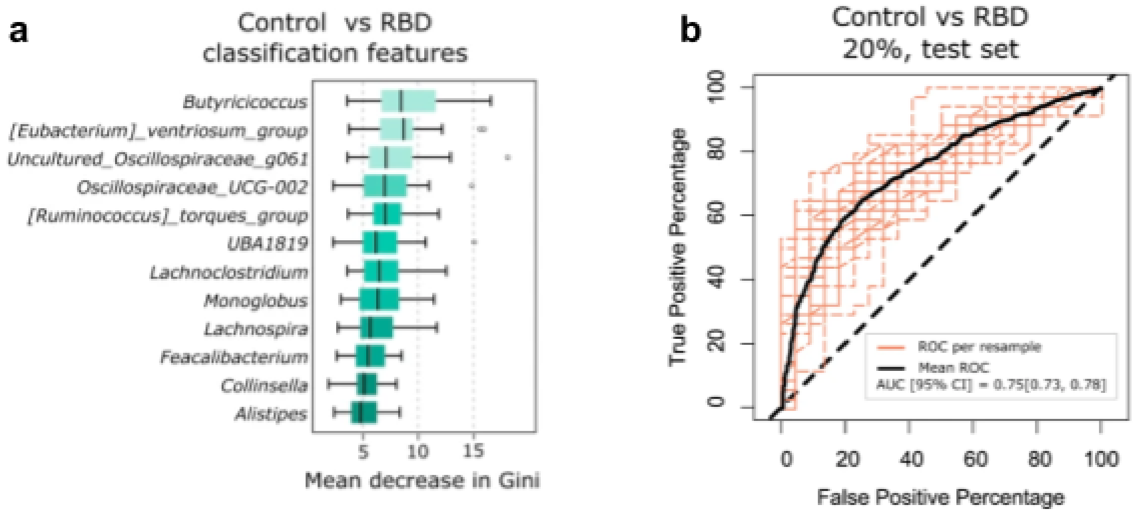


**Figure S1: Performance of the existing Random Forest Model for classification of RBD and control.** a) Control vs. RBD classification features selected through RFE. b) Receiver Operating Characteristic Curve for the binary classifier. Figure from Huang et al., 2023, licensed under Creative Commons Attribution 4.0 International License (Rightslink® by Copyright Clearance Center)

#### Selection of Datasets

- PubMed search with specific terms
- Apply inclusion criteria

#### Hyperparameter Tuning

- Twelve genera from Huang et al. (2023)
- Run GridSearchCV to optimize model parameters

#### Integration of Clinical and Taxonomic Features

- Combine six clinical with twelve taxonomic features
- Train and test predictive model

#### Meta-Analysis Workflow

1. Preprocessing
    a. Download reads from ENA Browser
    b. Perform quality check (FastQC + MultiQC) and control (fastp)
    c. Generate feature table via QIIME2
2. Batch Correction: Apply MBECS to remove batch effects
3. Differential Abundance Analysis: Use ANCOM-BC, adjust p-values by Benjamini-Hochberg method
4. Feature Selection
    a. Apply RFE, SFS-F, SFS-B, and SHAP.
    b. Average ranks across algorithms and select top ranked taxa
5. Model Building & Assessment
    a. Utilize Random Forest Classifier with 1000 bootstrap iterations
    b. Evaluate with AUC and additional metrics

**Figure S2: Methodology.** Flowchart of the methodology used for the study.

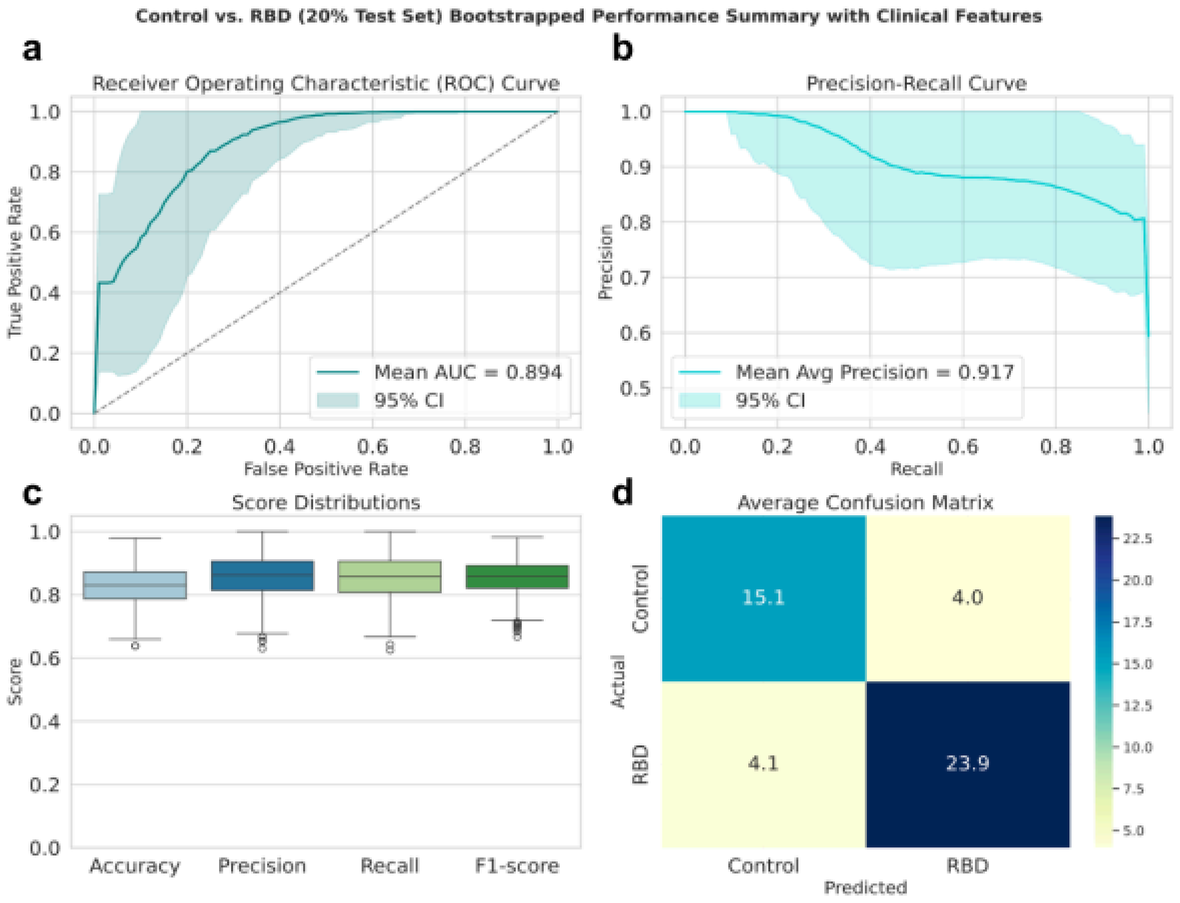


**Figure S3: Performance of the binary classifier with six clinical features.** Performance of the tuned binary classifier on the 20% test set using the six clinical features. a) The ROC curve showed a mean AUC of 0.894 on the test set across 1000 bootstrapping iterations. b) The Precision-Recall curve showed Mean Average Precision to be 0.917. c) The miscellaneous performance scores showed that the mean accuracy was 0.83, the mean precision was 0.87, the mean recall was 0.85, and the mean F1-score was 0.84. d) The confusion matrix.

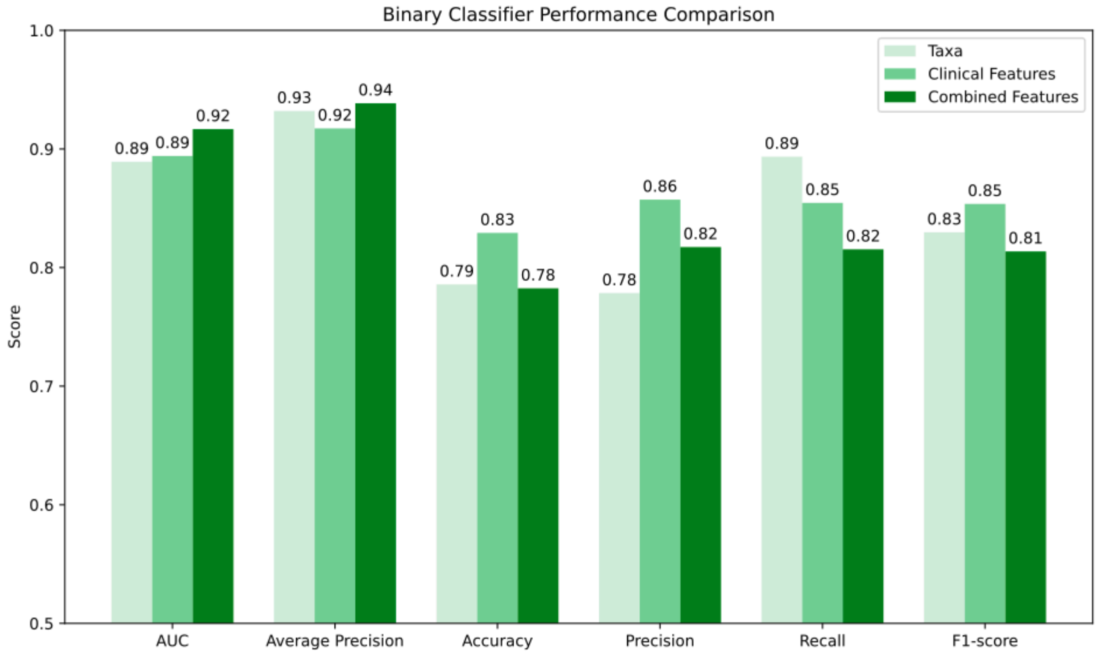


**Figure S4: Comparison of the model performance across taxonomic and clinical features**.

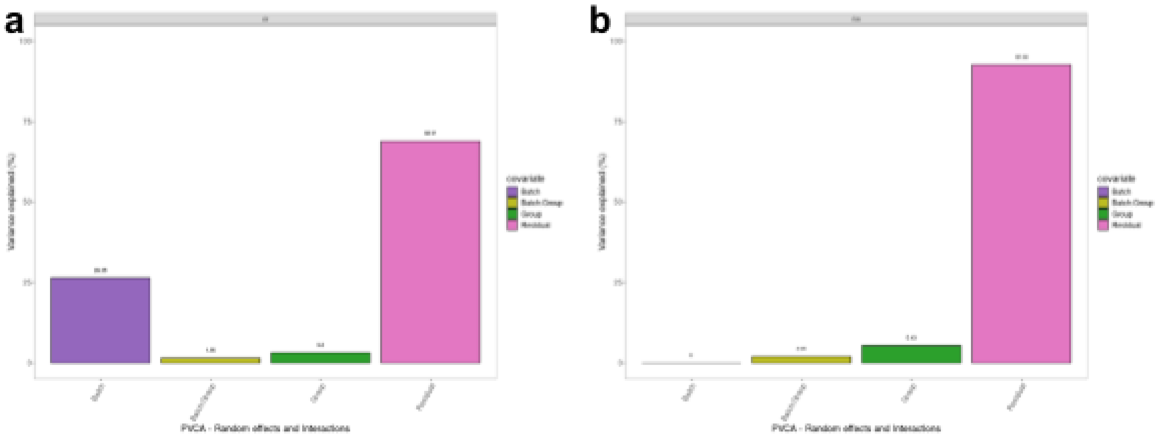


**Figure S5: Principal Variance Component Analysis (PVCA) plots**. a) before and b) after batch correction (rbe). After batch correction, the variation due to batch effects is reduced from 26% to 0%.

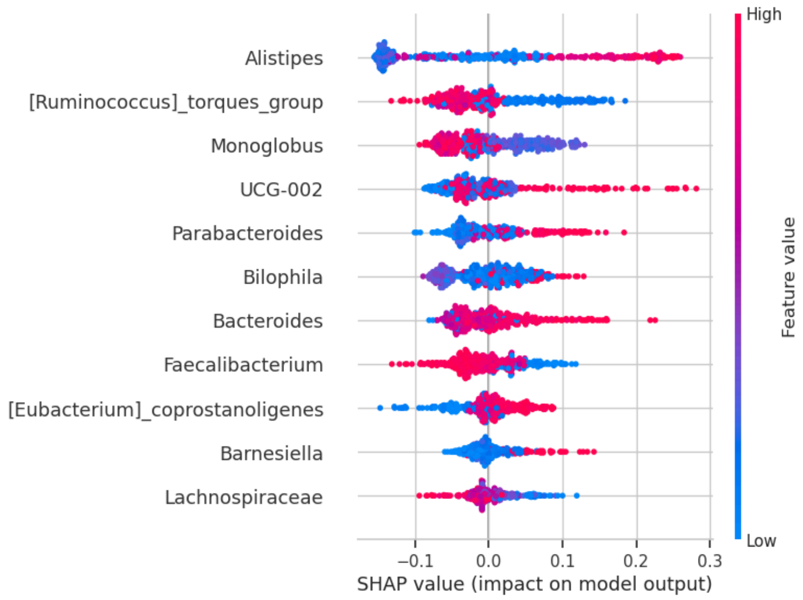


**Figure S6: SHAP Summary Plot for eleven top- ranked taxonomic features.**

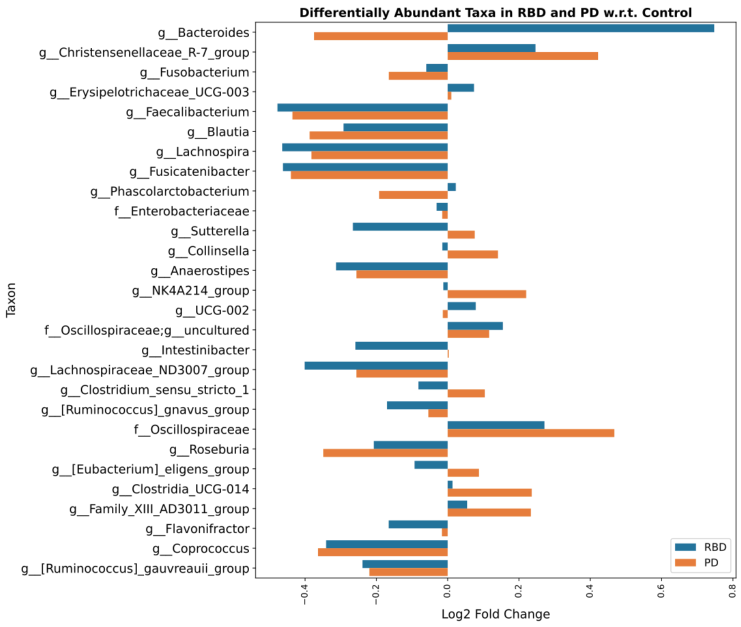


**Figure S7: Differentially abundant taxa in RBD and PD.**

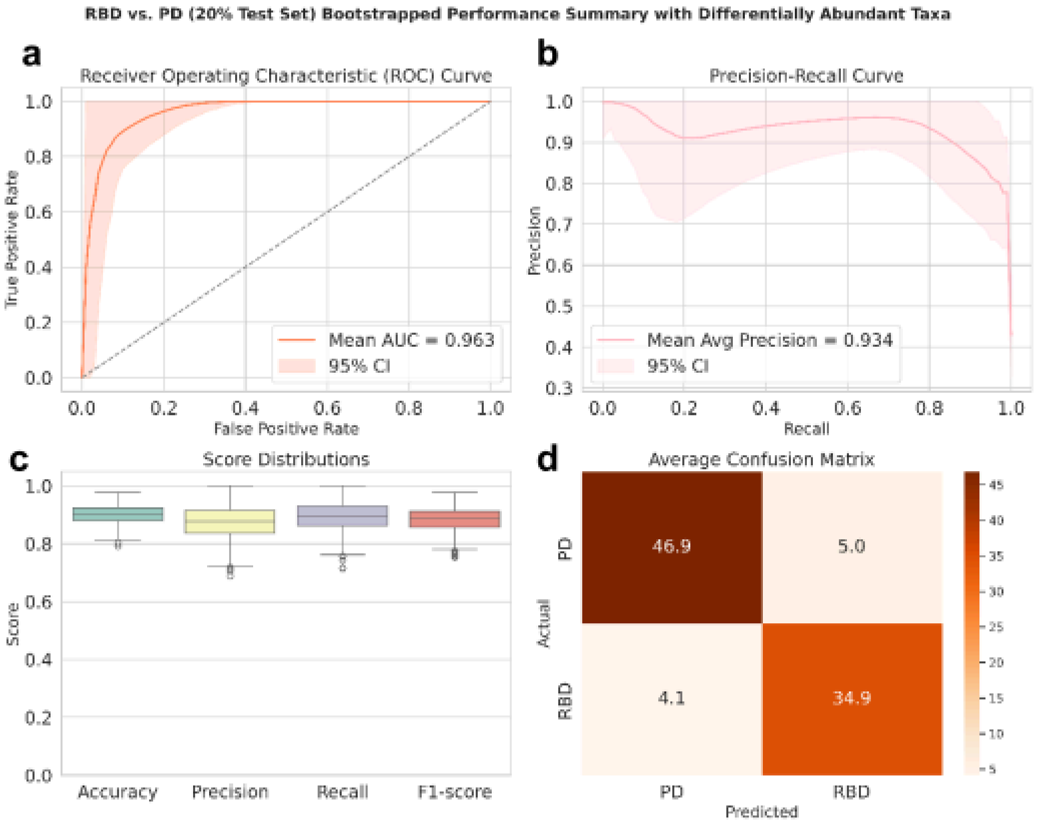


**Figure S8: Model Performance for RBD vs. PD**. Performance of the tuned binary classifier on the 20% test set using the differentially abundant taxa. a) The ROC curve showed a mean AUC of 0.963 on the test set across 1000 bootstrapping iterations. b) The Precision-Recall curve showed Mean Average Precision to be 0.934. c) The miscellaneous performance scores showed that the mean accuracy was 0.9, the mean precision was 0.87, the mean recall was 0.9, and the mean F1-score was 0.87. d) The confusion matrix.
